# Spatial Metabolomics by Desthiobiotin Ligase (DESTNI) in Live Cells

**DOI:** 10.64898/2026.07.05.736296

**Authors:** Chang-Mo Yoo, Jae-Yoon Jo, Chang-Ryul Choi, Ye Seop Park, Yun Jae Cha, Sungwoon Jung, Jiwoong Kang, Jeesoo Kim, Yun Pyo Kang, Tae Hyeon Yoo, Jong-Seo Kim, Hyun-Woo Rhee

## Abstract

Proximity labeling has transformed spatial proteomics by enabling compartment-resolved mapping of protein environments in living cells, yet its extension to small-molecule metabolites has not been demonstrated, probably due to limitations in labeling chemistry and labeled metabolites identification. Here, we introduce DESTNI, an engineered desthiobiotin (DTB) ligase derived from TurboID through directed evolution, and establish a platform for spatially resolved profiling of amine-containing metabolites. A directed evolution strategy based on a yeast display system yielded DESTNI with efficient DTB-dependent reactivity, enabling robust and compartment-specific proximity labeling across diverse subcellular environments. To identify the DTB-modified amino metabolome, we developed an integrated analytical framework combining DTB-modified amino metabolite standards, *in vitro* DESTNI profiling, and *in silico* MS/MS prediction, enabling systematic annotation of DTB-modified amino metabolites. To extend this chemistry to metabolites, we combined synthetic DTB-conjugated metabolite reference standards, *in vitro* DESTNI-reactive metabolite discovery, and machine-learning prediction of DTB-derivatized metabolites and oligopeptides. Organelle-targeted DESTNI recovered reproducible compartment-enriched amino metabolite signatures, including mitochondrial matrix-enriched glycine, 5-aminolevulinic acid, ornithine and spermidine adducts, as well as nuclear-enriched γ-aminobutyric acid and 5-aminovaleric acid adducts. Together, this work establishes DESTNI as a proximity labeling platform that bridges spatial proteomics and metabolomics and provides a general strategy for mapping subcellular biochemical environments in living cells.

## Introduction

The spatial organization of biomolecules is a defining feature of cellular identity and function. While recent advances in spatial proteomics have revealed compartment-resolved protein landscapes in living cells, spatial metabolomics remains substantially more limited. Imaging-based approaches are typically applied to fixed or processed samples and offer limited MS/MS-based metabolite identification, whereas fractionation-based strategies often suffer from metabolite leakage, low spatial fidelity, and incompatibility with dynamic measurements in intact cells^1^. These limitations underscore the need for approaches that can capture localized metabolic environments in living cells without physical separation.

Proximity labeling (PL) has emerged as a powerful strategy to map spatially restricted biomolecules by enzymatically generating reactive intermediates that covalently modify nearby targets. Among these tools, TurboID—an engineered biotin ligase with rapid kinetics—has enabled robust proximity proteomics in living systems^2^. Notably, the underlying AMP-mediated chemistry of TurboID suggests potential reactivity toward primary amine functionalities, motivating interest in extending proximity labeling strategies to small amino metabolites. However, extending TurboID-based labeling to metabolomics presents fundamental challenges. Biotin-modified amino metabolites exhibit inefficient recovery from streptavidin, limiting analytical coverage. In addition, TurboID-based labeling requires supplementation with exogenous biotin to achieve robust activity, which can elevate cellular biotinylation beyond physiological levels and potentially introduce metabolic or developmental perturbations^3^. Furthermore, in tissues with high endogenous biotinylation, detecting TurboID-labeled metabolites by mass spectrometry is severely confounded because both endogenous and engineered modifications generate the same mass adduct. Together, these features constrain the applicability of biotin-based proximity labeling strategies for spatial metabolomics.

These limitations of biotinylation-based strategies in spatial metabolomics highlight the need for an alternative substrate such as desthiobiotin (DTB) for proximity-based metabolite labeling. DTB is a biotin analog that retains high-affinity binding to streptavidin while enabling mild and efficient elution, thereby substantially improving recovery of labeled species and analytical coverage. This principle has been demonstrated in DTB-modified peptide identification workflows^4^, suggesting that DTB chemistry could also support small-molecule proximity labeling if paired with an enzyme capable of efficient DTB activation.

To harness the improved recovery and minimal interference offered by DTB, we developed DESTNI as a DTB-optimized engineered ligase that bridges proteomics and metabolomics in a unified framework. DESTNI retains the targeting versatility of TurboID while generating DTB-reactive intermediates compatible with both protein and metabolite labeling. Importantly, DTB-modified lysine residues (Lys +196 Da) are readily distinguished from endogenous biotinylation events (Lys +226 Da), enabling cleaner identification of enzymatically labeled protein products while preserving the chemical advantages needed for metabolite recovery.

Here we establish DESTNI as a platform for spatially resolved profiling of amine-containing metabolites in living cells. The study proceeds in four linked steps: engineering of a DTB-utilizing ligase, validation that DTB-dependent labeling remains spatially resolved after organelle targeting, construction of reference and machine-learning-based spectral libraries, and application of the resulting workflow to compartment-resolved metabolite profiling. In this framework, enzyme engineering provides spatial control, spectral libraries supply molecular identity, and organelle targeting connects labeled metabolites to subcellular context.

## Results

### Directed evolution produces a desthiobiotin-utilizing proximity-labeling enzyme

To develop a proximity-labeling enzyme capable of efficiently using desthiobiotin (DTB), we performed yeast surface display-based directed evolution starting from TurboID (**Fig. 1a**). The TurboID library of approximately ten million variants was generated by error-prone PCR, displayed on the yeast surface as Aga2p fusions, and screened for DTB-dependent labeling using streptavidin staining and fluorescence-activated cell sorting (FACS). Under these screening conditions, TurboID showed only minimal DTB reactivity (**Fig. 1b**), reflecting its intrinsic preference for biotin and indicating that improved DTB utilization would require active enzyme engineering rather than reliance on pre-existing activity.

**Fig. 1.**
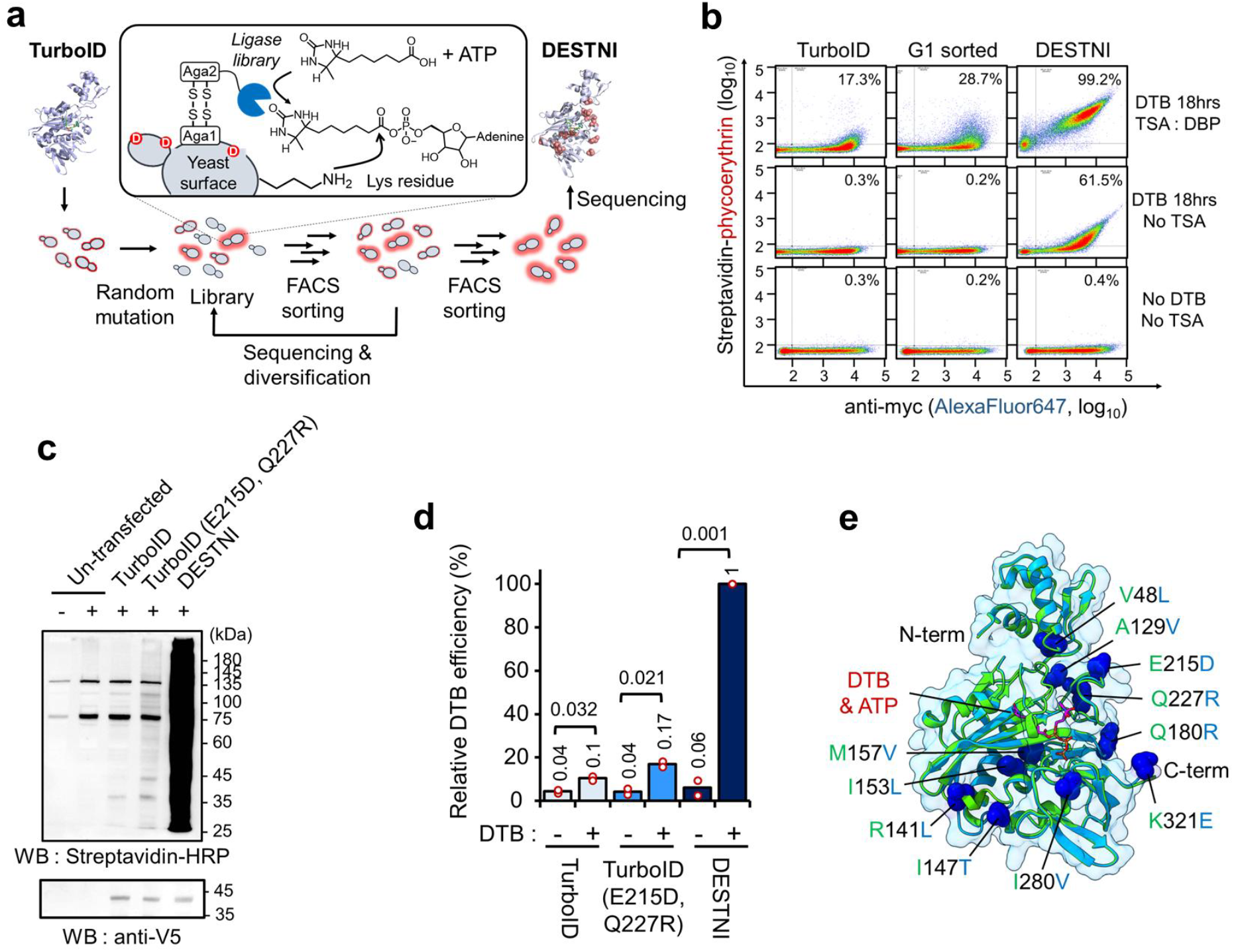
Directed evolution of DESTNI, a DTB-utilizing proximity-labeling enzyme. (a) Yeast surface display workflow used to evolve TurboID toward desthiobiotin (DTB) utilization. A randomized TurboID library was displayed as Aga2p fusions, incubated with DTB and sorted by FACS for DTB-dependent streptavidin labeling. (b) Representative flow-cytometry plots for TurboID, a first-generation enriched variant (G1-sorted) and DESTNI. Yeast cells were incubated with 50 μM DTB, 5 mM MgCl_2_ and 1 mM ATP for 18 h. Top panels include tyramide signal amplification (TSA), middle panels show DTB labeling without TSA and bottom panels show no-DTB controls. Streptavidin-phycoerythrin reports DTB labeling, and anti-Myc staining reports surface expression. Percentages indicate Q2/(Q2+Q4), where Q denotes flow cytometry quadrants. **(c)** Streptavidin-HRP blot analysis of DTB labeling by TurboID, the TurboID (E215D, Q227R) mutant and DESTNI expressed as Sec61B fusions in HEK293T cells. Cells were incubated with 500 μM DTB for 15 min. DTB-modified proteins were detected by streptavidin blotting, and ligase expression was assessed by anti-V5 immunoblotting. An overexposed streptavidin blot is shown to visualize weak labeling. **(d)** Quantification of DTB labeling from the experiment in (c). Streptavidin signal intensities were normalized to V5 expression. Individual biologically independent replicates (n = 2) are shown as red-edged white circles, and bars indicate mean values. **(e)** AlphaFold2-predicted structures of TurboID (light green) and DESTNI (light blue) shown as an aligned overlay. DTB (magenta) was positioned by *in silico* docking, and DESTNI substitutions (V48L, A129V, R141L, I147T, I153L, M157V, Q180R, E215D, Q227R, I280V and K321E) are shown as blue spheres.

In the first stage of directed evolution, six rounds of screening were carried out while progressively increasing stringency. Early rounds used long DTB incubations (20 hours) at 50 μM together with a desthiobiotin-phenol (DBP)-mediated tyramide signal amplification (TSA) step2 to enable detection of extremely low activities (**Supplementary Fig. 1**). The labeling duration was gradually reduced across rounds, and in the sixth round, variants were enriched after only a 6-hour DTB incubation with the TSA step. This strategy yielded a population with measurably improved DTB reactivity (**Fig. 1b**).

The enriched first-generation pool (G1-sorted) was further diversified by error-prone PCR to generate a second-generation library, which was subjected to four rounds of sorting under stricter conditions. The contribution of TSA was gradually reduced by lowering the concentration of DBP, and the final round was performed entirely without TSA. DTB incubation was kept at 50 μM for 6 hours throughout these rounds to maintain a consistent selection pressure. Through this two-stage directed evolution campaign, we isolated an engineered enzyme, which we named DESTNI (**Fig. 1b, Supplementary Fig. 1**).

Sequencing revealed eleven amino acid substitutions that distinguish DESTNI from TurboID: V48L, A129V, R141L, I147T, I153L, M157V, Q180R, E215D, Q227R, I280V, and K321E. To determine whether the activity observed in the yeast display system was retained in mammalian cells, we compared TurboID, the TurboID (E215D, Q227R) mutant, an intermediate variant identified from the G1-sorted population, and DESTNI as Sec61B fusions in the ER lumen of HEK293T cells and labeled the cells with DTB for 15 min. DESTNI generated strong DTB-dependent protein labeling, whereas TurboID and the TurboID (E215D, Q227R) mutant showed only weak activity under the same conditions (**Fig. 1c,d**). Parallel probe-comparison experiments further confirmed robust DTB-labeling activity of DESTNI in mammalian cells (**Supplementary Fig. 2**).

Mapping the DESTNI substitutions onto an AlphaFold2-predicted structure^5^ indicated that most substitutions are not directly involved in first-shell contacts for the modeled DTB/ATP adenylation chemistry. Instead, several substitutions, including A129V, R141L, I153L, and M157V, form a second-shell network around the adenylation pocket, whereas V48L and I280V are positioned more distally (**Fig. 1e**). Among these sites, the M157 residue is especially notable because the Ellington group previously implicated mutations at this position in altered substrate preference of BirA variants engineered toward DTB utilization^6^. This arrangement suggests that this second-shell remodeling tunes pocket geometry or dynamics to facilitate productive DTB binding and ATP-dependent positioning for adenylation, rather than creating a single new catalytic contact.

Functional analysis of DESTNI single-reversion mutants supported this structure-guided interpretation. In a cytosolic DESTNI-V5-NES background, reverting individual DESTNI-specific residues to their TurboID counterparts showed that V129A, L141R, L153I, V157M, D215E and R227Q significantly reduced V5-normalized DTB labeling relative to DESTNI (**Supplementary Fig. 2**). L153I was among the strongest single-reversion losses, and V157M provided functional support for the importance of the M157 position previously implicated by Ellington and colleagues. By contrast, reversions at L48, T147, R180, V280 and E321 in DESTNI had less pronounced effects on DTB-labeling activity, indicating that these positions make smaller individual contributions to the enhanced DTB activity of DESTNI. These results indicate that the high activity of DESTNI toward DTB is generated by synergistic contributions from mutations distributed in the second shell. The success of engineering DESTNI demonstrates that iterative yeast-display screening yielded a DTB-optimized proximity-labeling enzyme that functions robustly in living cells.

### Organelle-targeted DESTNI establishes spatially resolved DTB labeling in cells

Having evolved DESTNI for DTB utilization, we next asked whether the new substrate preference preserved the spatial precision that makes proximity labeling useful. DESTNI was targeted to the mitochondrial matrix (MTS), outer mitochondrial membrane (TOM20), cytosol (NES), nucleus (NLS), endoplasmic reticulum membrane (Sec61B), peroxisomes (PEX3), plasma membrane (CAAX) and lysosomes (TMEM192). Organelle-targeted DESTNI constructs were expressed in HEK293T cells and incubated with 500 μM DTB for 15 min. Immunofluorescence imaging revealed that DTB-labeled species, detected by streptavidin-conjugated fluorophore or HRP, co-localized spatially with DESTNI expression, confirming that the evolved enzyme could be redirected to distinct subcellular environments (**Fig. 2a,b, Supplementary Fig. 3**).

**Fig. 2.**
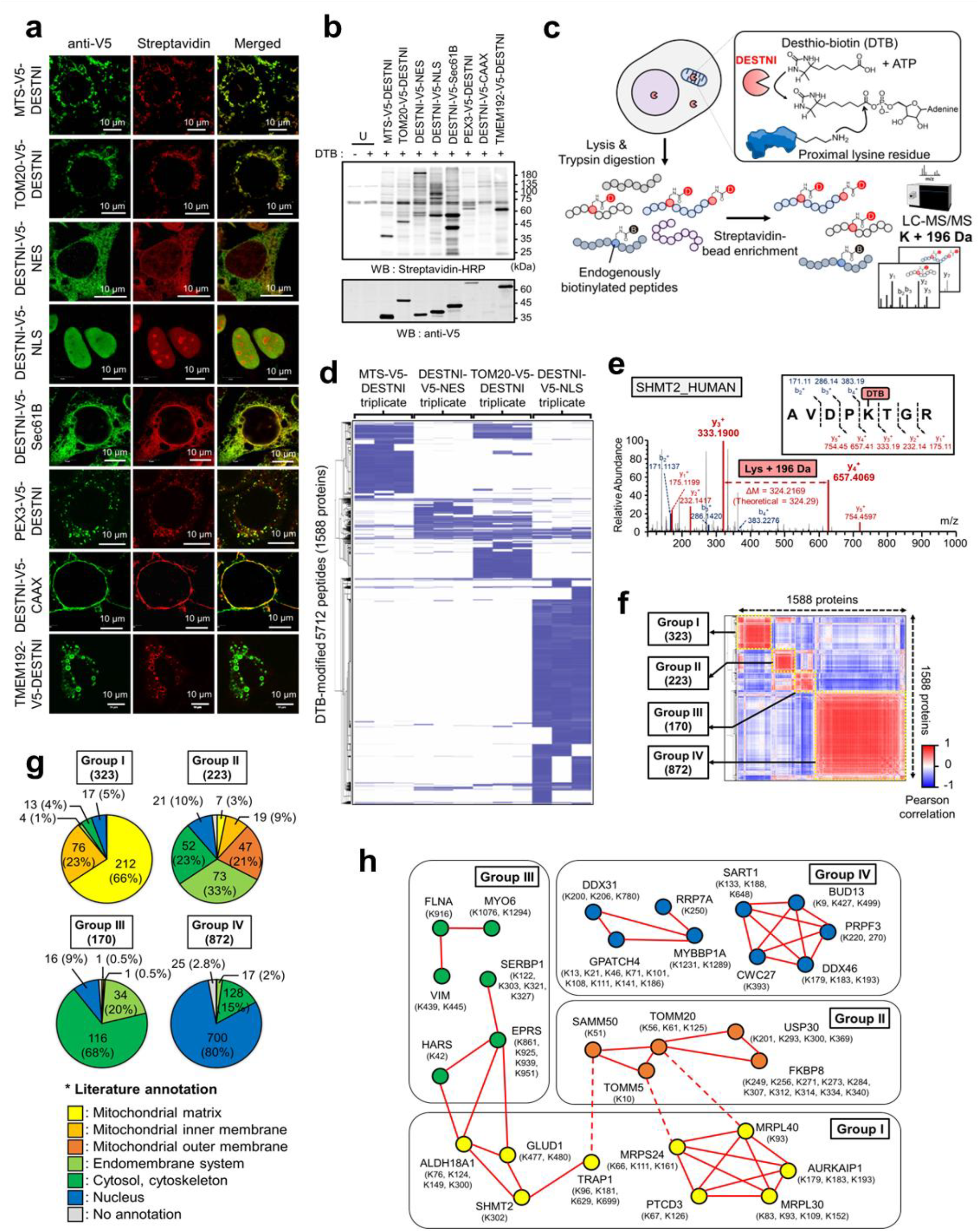
Organelle-targeted DESTNI enables compartment-specific proximity-labeling proteomics. **(a)** Immunofluorescence imaging of organelle-targeted DESTNI constructs expressed in HEK293T cells. Cells were treated with 500 μM DTB for 15 min. DTB-labeled species were visualized with streptavidin-AF568, and DESTNI expression was detected with anti-V5 antibody followed by AF488-conjugated secondary antibody. Targeting sequences directed DESTNI to the mitochondrial matrix (MTS), outer mitochondrial membrane (TOM20), cytosol (NES), nucleus (NLS), endoplasmic reticulum membrane (Sec61B), peroxisomes (PEX3), plasma membrane (CAAX) or lysosomes (TMEM192). Scale bars, 10 μm. **(b)** Immunoblot analysis of organelle-targeted DESTNI variants. DTB-labeled proteins were detected by streptavidin-HRP, and DESTNI expression was detected by anti-V5 immunoblotting. U denotes untransfected control cells without (lane 1) or with (lane 2) DTB treatment. **(c)** Workflow for enrichment and LC–MS/MS analysis of DTB-modified peptides generated by organelle-targeted DESTNI. **(d)** Peptide-level heatmap of DTB-modified peptides (5,712 peptides; 1,588 proteins) across four organelle-targeted DESTNI constructs (MTS, TOM20, NES and NLS; n = 3). Intensities were log_2_-transformed and globally normalized by cyclic loess; rows were clustered using one-minus Pearson correlation distance and average linkage. **(e)** Representative MS/MS spectrum of the DTB-modified SHMT2 peptide AVDPK(DTB)TGR, showing characteristic DTB-specific fragment ions. **(f)** Pearson correlation matrix calculated from cyclic-loess-normalized protein intensities for 1,588 proteins identified from DTB-modified peptides. Correlation-based clustering resolved four major protein groups (Groups I–IV). **(g)** Subcellular localization annotations for the protein groups defined in (f). Pie charts show the distribution of annotated localizations within each group. **(h)** STRING network of representative DESTNI-labeled spatial proteins. Nodes are color-coded by subcellular compartment, and numbers in parentheses indicate the number of DTB-modified lysines identified per protein. Red lines indicate STRING protein-protein associations.

Having established compartment-specific labeling by imaging and immunoblotting, we next examined the specificity of DESTNI-mediated proximity labeling using mass spectrometry. We focused on four representative constructs—MTS-DESTNI, TOM20-DESTNI, DESTNI-NES, and DESTNI-NLS—and enriched DTB-modified peptides (Lys +196 Da) for LC-MS/MS analysis (**Fig. 2c**). DTB-modified peptides derived from these experiments mapped to proteins associated with the corresponding subcellular compartments, yielding distinct labeling profiles for each DESTNI localization. Consistent with this, a hierarchically clustered heatmap of peptide-level labeling patterns revealed clear separation according to subcellular targeting (**Fig. 2d**). A representative MS/MS spectrum of a DTB-modified peptide from the mitochondrial matrix enzyme SHMT2 illustrates the characteristic DTB-specific fragmentation pattern used for confident peptide identification (**Fig. 2e**).

To further compare labeling relationships across compartments, we computed a Pearson correlation matrix based on cyclic-loess-normalized DTB-labeled intensities for 1,588 proteins (**Fig. 2f**). Correlation-based clustering resolved four major groups of proteins with shared spatial labeling profiles (Groups I–IV). Functional annotation of these groups revealed enrichment for proteins assigned to the mitochondrial matrix, outer mitochondrial membrane, cytosol or nucleus, respectively, consistent with the four DESTNI targeting sites (**Fig. 2g**). To illustrate representative examples, we visualized consistently DTB-labeled proteins in a subcellular interaction network, with nodes color-coded by compartment and edges representing protein-protein interactions obtained using default STRING parameters^7^ (**Fig. 2h**). These data established that DTB labeling by DESTNI remains genetically targetable and spatially resolved in living cells, providing the necessary foundation for applying the same enzyme to metabolite capture.

### Building a hybrid spectral-library framework for DESTNI metabolomics

For the metabolomics applications of DESTNI, we assessed the compatibility of DESTNI-mediated labeling with streptavidin-based metabolite enrichment and found that DTB-modified metabolites generated by DESTNI were efficiently recovered in elution fractions, in contrast to biotin-labeled metabolites produced by TurboID, which showed limited recovery (**Supplementary Fig. 4**). This result established DTB as a practical enrichment handle for small-molecule labeling, while leaving metabolite identities to be resolved by LC-MS/MS acquisition and spectral-library matching. We then validated the core chemistry used by the metabolomics workflow. Purified DESTNI generated the DTB-AMP intermediate in vitro, whereas boiled enzyme controls showed minimal signal, confirming enzyme-dependent adenylation of DTB (**Fig. 3a,b, Supplementary Fig. 5**). In a targeted *in vitro* reaction, DESTNI labeled taurine to generate DTB-taurine with the expected precursor and MS/MS fragments (**Fig. 3c**). Because these products were expected to be low-abundance and to fragment through a shared DTB headgroup, we established a capillary LC-MS/MS workflow using an in-house packed 150 μm i.d. C18 column and stepped HCD collision energies. These conditions balanced sensitivity and chromatographic reproducibility (**Fig. 3d, Supplementary Figs. 6 and 7**).

**Fig. 3.**
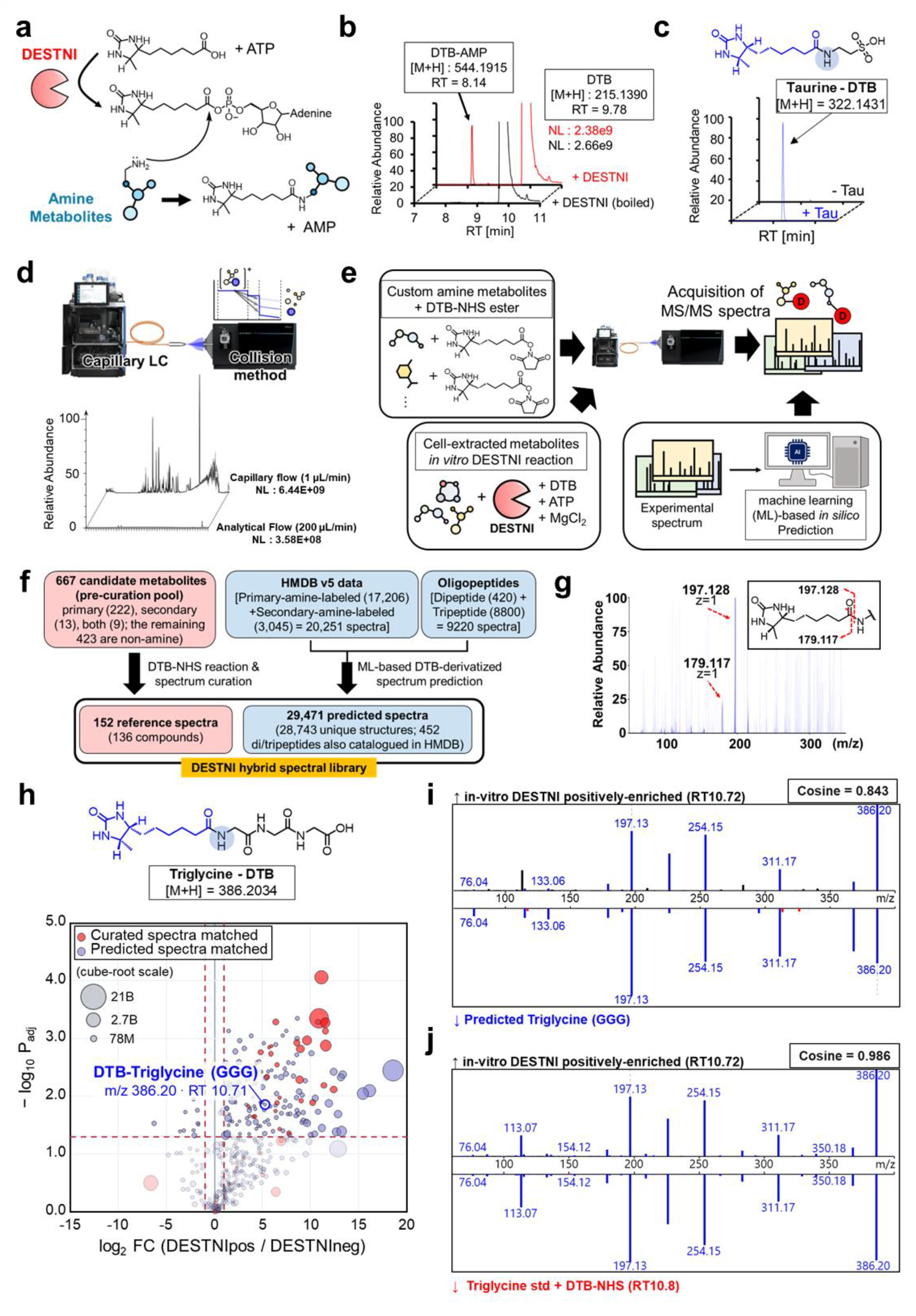
DESTNI-mediated labeling and DTB-modified spectral-library construction for amino metabolites. **(a)** DESTNI-mediated labeling mechanism for amino metabolites. **(b)** Detection of DTB-AMP generated by purified His10-MBP-DESTNI *in vitro*. Reactions containing DTB and ATP were analyzed by LC–MS/MS, and extracted ion chromatograms (XICs) for DTB-AMP ([M+H]+ = 544.1915, RT = 8.14 min) are shown. Boiled DESTNI served as a negative control. **(c)** *In vitro* labeling of taurine by DESTNI. Reactions were performed with or without taurine and analyzed by LC–MS/MS. Representative XICs and an MS/MS spectrum of DTB-taurine ([M+H]+ = 322.1431) are shown. **(d)** Capillary LC–MS/MS optimization workflow for DTB-modified metabolites. **(e)** Workflow for construction of the hybrid DTB-amine spectral library from DTB–NHS-derivatized metabolite standards, *in vitro* DESTNI reactions using HEK293 T-REx metabolite extracts and ML-based MS/MS spectrum prediction. **(f)** Composition of the hybrid spectral library of DTB-modified amino metabolites integrating reference experimental spectra with ML-based predicted spectra for HMDB amine-containing metabolites and oligopeptides. **(g)** Overlaid MS/MS spectra of DTB-modified standards, highlighting characteristic DTB headgroup fragmentation and diagnostic fragment ions at m/z 197.128 and 179.117. **(h)** Volcano plot of DTB-modified metabolite features identified from *in vitro* DESTNI reactions by matching to the hybrid DTB-amine spectral library. Features are plotted as log_2_ fold change (DESTNI-positive/DESTNI-negative; n = 3 biologically independent samples per group) versus -log_10_ P value (two-sided Welch’s t-test). DTB-triglycine is highlighted. **(i)** Mirror plot comparing the experimental MS/MS spectrum acquired from the *in vitro* DESTNI reaction (top) with the corresponding ML-predicted spectrum (bottom) for DTB-triglycine (cosine similarity = 0.843). **(j)** Mirror plot comparing the experimental MS/MS spectrum acquired from the *in vitro* DESTNI reaction (top) with the MS/MS spectrum of DTB-NHS-derivatized triglycine standard (bottom) (cosine similarity = 0.986), confirming the DTB-triglycine assignment.

We next established the metabolomics workflow to address analytical problem that limits proximity-based spatial metabolomics. It is noteworthy that metabolomics analysis setup is quite different from that of proteomics. The striking difference is that metabolite detection requires “standard” molecules or reference spectral library to confirm the detection (e.g. cosine value calculation of MS/MS spectra) while peptide MS/MS spectra can be assigned by database searching against predictable peptide sequences^8^. Because public metabolite spectral libraries provide limited coverage of DTB-modified metabolites, we need to chemically generate reference spectral library of DTB-modified metabolites for the confirmation of the DTB-modified metabolites using DESTNI.

First, we constructed a reference spectral library of DTB-modified amino metabolites rather than relying on public databases. The experimental reference arm used DTB–NHS derivatization and LC-MS/MS analysis of a 667-compound candidate metabolite panel. This panel comprised 222 primary-amine candidates, 13 secondary-amine candidates, 9 compounds containing both primary and secondary amines, and 423 non-amine candidates. Manual validation of the resulting spectra yielded 152 high-confidence DTB-modified spectra representing 136 compounds (121 primary-amine, 9 secondary-amine, 6 compounds containing both primary and secondary amines), including di-DTB species and chromatographically resolved regioisomers where applicable (**Fig. 3e,f, Supplementary Fig. 7**). Across this reference set, the DTB headgroup fragments described above served as a shared chemical signature for recognizing DTB-modified amino metabolites, with DTB-specific diagnostic ions at m/z 197.128 and 179.117 reproducibly observed across the library (**Fig. 3g**).

To expand beyond measured reference spectra, we generated a predicted DTB-derivatized spectral library containing DTB diagnostic ions (m/z 197.128). We computationally generated all possible DTB-labeled structures for HMDB v5 metabolites^9^ using the RDKit software^10^, targeting primary-amine sites (17,206 spectra) and secondary-amine sites (3,045 spectra), yielding 20,251 HMDB-derived predicted spectra. In parallel, all possible DTB-labeled dipeptides (420 spectra) and tripeptides (8,800 spectra), in which DTB was conjugated to either the lysine residue or N-terminus of the peptides, were generated computationally to include short oligopeptide-like metabolites. A machine-learning spectral prediction model trained on the 152 experimentally measured reference spectra was then used to predict 29,471 spectra in total, corresponding to 28,743 unique structures; 452 structures overlapped between the HMDB-derived and oligopeptide spaces. Together, the reference and predicted libraries constituted a hybrid spectral library (**Fig. 3f**).

Finally, we asked whether this framework captured products formed by DESTNI in a cellular metabolite background. Triplicate *in vitro* reactions with HEK293 T-REx metabolite extracts yielded 5,619 DTB-candidate features containing DTB diagnostic ions at m/z 197.128. Among these, 1,520 showed statistically significant enrichment in DESTNI-positive reactions relative to no-enzyme controls, using a log_2_ fold change > 1 and a Benjamini-Hochberg-adjusted P < 0.05 as thresholds. The hybrid library matching annotated 146 enriched features: 30 with measured reference DTB-conjugate spectra and 116 with predicted spectra. (**Fig. 3h, Supplementary Fig. 8**).

Among the DESTNI-modified metabolites in the extract, a feature annotated as DTB-triglycine matched the spectrum predicted by the machine-learning model (cosine similarity = 0.843) (**Fig. 3i**). We validated this feature by chemical preparation of DTB-triglycine through DTB-NHS derivatization of triglycine standard compound. It produced a stronger correlative MS/MS spectrum with the feature detected in the *in vitro* DESTNI reaction (cosine similarity = 0.986) (**Fig. 3j**). Thus, we confirmed that our hybrid spectral library was useful for annotating DTB-modified metabolites.

### Organelle-targeted DESTNI captures compartment-enriched amino metabolite signatures

Having established a metabolomics analysis workflow using DESTNI, we asked whether localized DESTNI activity could report spatial metabolite information in intact cells. Stable HEK293 T-REx cell lines expressing mitochondrial matrix-targeted DESTNI (MTS-DESTNI), cytosolic DESTNI (DESTNI-NES) or nuclear DESTNI (DESTNI-NLS) were incubated with DTB (500 μM) for 3 h. After labeling, metabolites were extracted with 80% MeOH, reconstituted in aqueous buffer, enriched on streptavidin beads, eluted with acidified 80% ACN, dried, reconstituted with an internal standard and analyzed by LC-MS/MS (Lumos). DTB-candidate MS/MS spectra were extracted from mzML files using diagnostic reporter-ion detection, spectral purity filtering and precursor-mass constraints before matching to reference and predicted DTB spectral libraries. No-probe and no-enzyme control samples were also processed using the same workflow to define background DTB-modified signals (**Fig. 4a**).

**Fig. 4.**
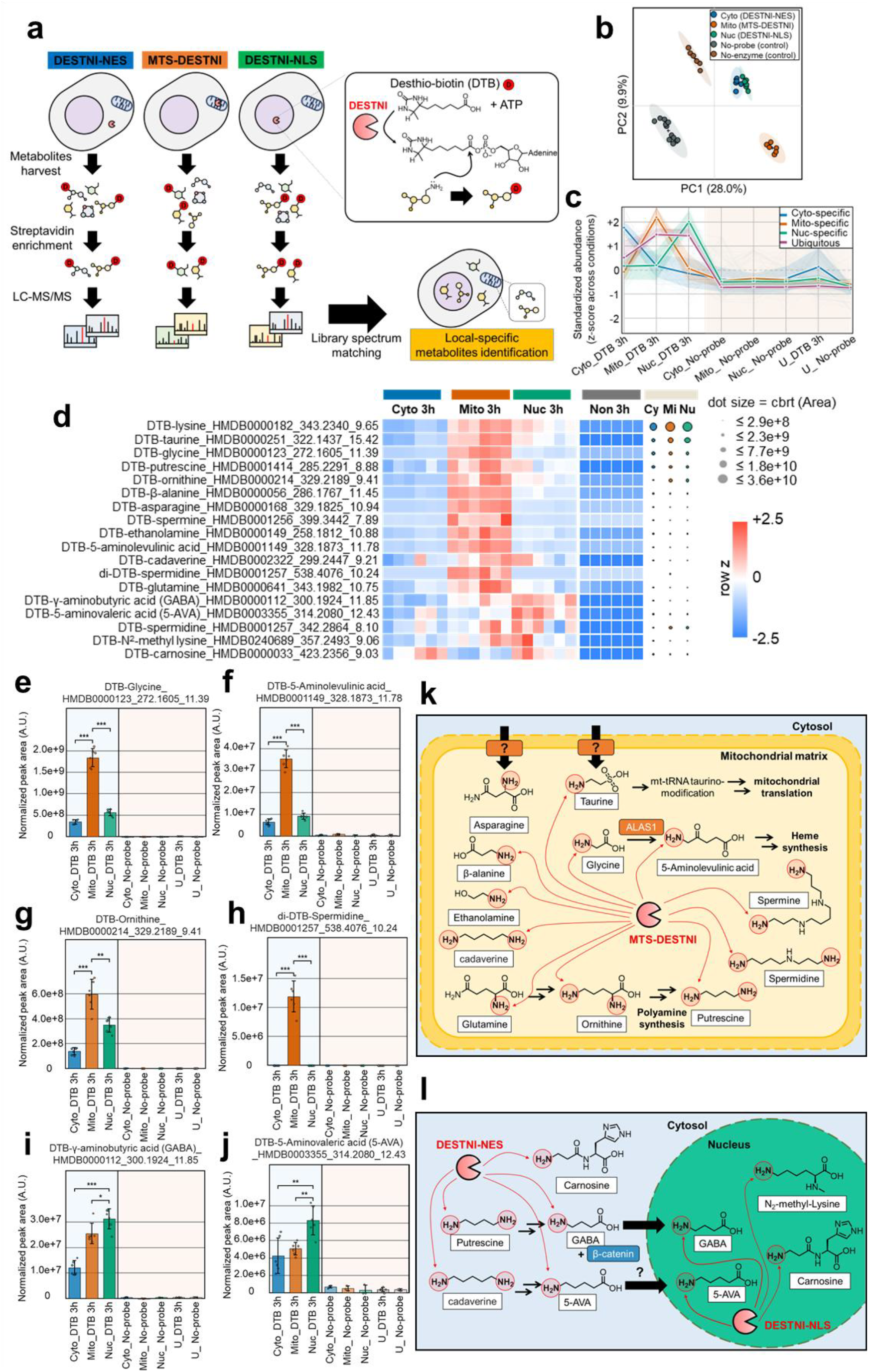
Organelle-targeted DESTNI enables spatial profiling of amino metabolites. **(a)** Spatial metabolomics workflow with DESTNI. Stable HEK293 T-REx cells expressing MTS-DESTNI, DESTNI-NES or DESTNI-NLS were labeled with DTB (500 μM) for 3 h, followed by metabolite extraction, streptavidin enrichment, LC–MS/MS analysis and DTB spectral-library matching. No-probe and no-enzyme controls were included to define background signals. **(b)** PCA plot of DTB-labeled metabolites identified across subcellular compartments. Biological replicates segregate according to DESTNI localization; ellipses show 95% normal-distribution confidence regions (n = 6 per group). **(c)** Ordering of DTB-labeled features by dominant compartmental signal. Features were grouped according to the condition with the highest internal-standard-normalized intensity across MTS-, NES- and NLS-DESTNI samples and negative controls. Rows were z-score scaled for visualization, highlighting mitochondria-, cytosol- and nucleus-dominant patterns as well as broadly distributed features. **(d)** Heatmap of reference-library-matched metabolites. Relative metabolite distributions are row-wise z-score normalized across all samples. Adjacent dot plots show cube-root-transformed, internal-standard-normalized peak areas; dot size is proportional to metabolite abundance. **(e-j)** Normalized LC–MS/MS peak areas for representative DTB-modified metabolites across organelle-targeted DESTNI samples: **(e)** DTB-glycine, **(f)** DTB-5-aminolevulinic acid, **(g)** DTB-ornithine, **(h)** di-DTB-spermidine, **(i)** DTB-GABA (γ-aminobutyric acid) and **(j)** DTB-5-aminovaleric acid. Bars indicate mean internal-standard-normalized peak areas; dots indicate biological replicates; error bars, s.d. (n = 6 per group). **(k)** Mitochondrial matrix-enriched DTB-labeled metabolites mapped to known mitochondrial metabolic pathways. Red arrows indicate metabolites detected by MTS-DESTNI; question marks denote proposed or untested steps. **(l)** Cytosol- and nucleus-enriched labeled metabolites. Red arrows indicate metabolites detected by DESTNI-NES or DESTNI-NLS; dashed arrows and question marks denote proposed or untested steps.

In principal component analysis (PCA), the resulting metabolite profiles separated according to DESTNI localization, indicating that each targeted enzyme sampled a reproducible local chemical environment (**Fig. 4b**). For visualization, DTB-labeled features were classified and ordered according to the subcellular targeting spaces for DESTNI (**Fig. 4c**). This ordering highlighted mitochondria-, cytosol- and nucleus-dominant patterns while preserving broadly distributed features. Reference-library matches and internal-standard-normalized dot plots showed that these patterns reflected coherent abundance differences across biological replicates (**Fig. 4d**).

The compartment-enriched profiles captured by reference-library matches were consistent with established subcellular biochemistry, providing clear biochemical validation for the approach. For instance, the strong DTB-taurine signal observed in the mitochondrial matrix aligns with the established role of taurine in forming modified wobble uridines in mitochondrial tRNAs^11^. MTS-DESTNI also preferentially detected specific subsets of expected amino acids and related precursors (**Fig. 4e–h,k**). The prominent signals for DTB-glycine and DTB-5-aminolevulinic acid are consistent with known mitochondrial metabolism: mitochondrial 5-aminolevulinic acid synthase condenses glycine with succinyl-CoA in the first committed step of heme biosynthesis^12^, whereas glycine also contributes to mitochondrial one-carbon metabolism^13^. Similarly, the detection of DTB-glutamine and DTB-ornithine reflects mitochondrial glutamate and ornithine metabolism centered on ornithine aminotransferase^14^.

Furthermore, MTS-DESTNI detected matrix-localized pools of ornithine-derived polyamines, including DTB-putrescine, DTB-spermine, and notably, di-DTB-spermidine. This robust enrichment underscores specialized polyamine biology within the mitochondria^15^. Given recent evidence that mitochondrial spermidine pools directly influence fatty acid oxidation machinery such as the mitochondrial trifunctional protein^16^, our findings provide further biochemical evidence supporting an active role for polyamines in the mitochondrial matrix. Beyond these expected features, MTS-DESTNI also enriched DTB-lysine, DTB-asparagine, DTB-β-alanine, DTB-ethanolamine and DTB-cadaverine, which were less directly predicted from canonical mitochondrial metabolism. These unexpected mitochondrial-enriched candidates suggest that localized DESTNI can capture additional amino acid, amino alcohol and diamine pools that become accessible to the mitochondria-targeted enzyme through local transport, exchange or organelle-localized metabolite availability.

NES- and NLS-targeted DESTNI resolved distinct patterns of spatial metabolite accessibility in the cytosol and nucleus (**Fig. 4i,j,l**). The most prominent pattern emerged from DESTNI-NLS, which preferentially recovered DTB-GABA and DTB-5-aminovaleric acid, together with additional enrichment of DTB-N^2^-methyl lysine and DTB-carnosine. DESTNI-NES provided a complementary cytosol-localized profile, in which DTB-carnosine was also detected, suggesting that this histidine-containing dipeptide may be accessible across multiple labeling environments. Collectively, these metabolite pools, particularly those represented by GABA, 5-aminovaleric acid, N^2^-methyl lysine and carnosine, warrant future mechanistic analysis. Taken together, we validated that DESTNI can reveal previously inaccessible spatial layers of the metabolome in live cells.

## Discussion

DESTNI demonstrates that proximity-labeling chemistry can be repurposed from spatial proteomics toward spatial metabolomics when enzyme engineering, spectral annotation and genetic targeting are developed together. A central advantage of the approach lies in the versatility of the desthiobiotin (DTB) tag. Within the same workflow, DTB serves as the substrate used by DESTNI to label proximal amine-containing metabolites, as a direct streptavidin-enrichment handle (K_D_ ≈ 10^-13^ M)^17^ that avoids additional click-chemistry reactions while retaining mild elution conditions, and as a mass-spectrometric barcode. Upon MS/MS fragmentation, the DTB moiety reliably generates dominant diagnostic reporter ions at m/z 197.128 and 179.117, facilitating recognition of DTB-modified metabolites.

Beyond these expected biochemical anchors, spatial profiling revealed localized accumulation of unexpectedly enriched metabolites whose canonical biosynthetic pathways typically reside elsewhere, highlighting unexplored local synthesis mechanisms or uncharacterized subcellular transporters. For instance, although asparagine is synthesized de novo in the cytosol by asparagine synthetase from aspartate and glutamine^18, 19^, its robust detection by MTS-DESTNI suggests either an unknown mitochondrial import mechanism or uncharacterized mitochondrial synthesis. Similarly, nuclear-enriched GABA and 5-aminovaleric acid raise interesting questions regarding their import and functions in the nucleus. Interestingly, GABA has emerging non-synaptic, gene-regulatory roles^20^, including the stabilization of nuclear β-catenin to activate Wnt/β-catenin signaling in cancer^21^. Overall, our findings offer biochemical evidence consistent with nuclear enrichment of GABA and 5-aminovaleric acid, supporting its evolving roles in the nucleus.

The spatially resolved detection of dipeptides and tripeptides, including those observed specifically in the mitochondrial matrix or nucleus (**Supplementary Fig. 9**) also suggests their specific subcellular accumulation could reflect localized activity of resident peptidases. Consequently, DESTNI provides a technical opportunity to map local peptide-metabolite generation and to investigate organelle-specific peptidase functions in living cells. The *in vitro* validation of DTB-triglycine illustrates this opportunity while also emphasizing that peptide-like assignments will require broader standard coverage and independent structural confirmation.

The concurrent detection of DTB-carnosine features is also chemically plausible because carnosine and related dipeptides are established nucleophilic quenchers of reactive carbonyl species^22^ that plays a beneficial role for healthspan. Our recent work also showed that taurine, spermidine and ethanolamine, can react with acyl-CoAs and biotinyl-AMP/acyl-AMP-like intermediates, thereby scavenging reactive acyl species (RAS)-mediated carbon stress and limiting aberrant protein acylation^23^. Because DESTNI generates the electrophilic DTB-AMP ester intermediate that can be categorized as RAS, DTB-modified carnosine together with taurine, ethanolamine, and spermidine may represent nucleophilic amine metabolite pools that are accessible to RAS in each compartment. Whether these metabolites undergo reactions with local pools of acyl-CoAs under carbon-stress conditions, and whether such reactions are protective for the proteostasis in the corresponding organelles, remains to be interesting questions for metabolite-driven protein modification fields^24^.

Despite these advances, certain limitations and future opportunities remain. At present, the DESTNI workflow targets amine-containing metabolites and therefore does not capture metabolite classes lacking suitable nucleophilic groups. A major bottleneck remains the annotation of DTB-diagnostic spectra. Within our current hybrid library framework, only 146 of the 1,520 statistically enriched DESTNI-reactive features (based on DTB diagnostic ions, 9.6%) could be assigned molecular identities. The remaining 90.4% (1,374 features) were left unannotated (**Supplementary Fig. 8**)—a substantial proportion of unidentified signals that mirrors the challenge of the “dark metabolome”^25^. In this context, our DESTNI-derived metabolomics datasets, which contain highly specific DTB diagnostic fragment signatures, can serve as a foundational resource to train future *in silico* metabolite-identification algorithms and help illuminate the unannotated spatial metabolome^26-28^. Overall, DESTNI establishes a genetically encoded enzymatic platform for spatially resolved metabolite labeling, providing a robust framework for the systematic interrogation of subcellular metabolic environments in living cells.

## Data availability

Data supporting the findings presented in this preprint are available within the article. Additional datasets generated and/or analyzed during the current study are available from the corresponding authors upon reasonable request.

## Acknowledgments

This work was supported by the Samsung Science and Technology Foundation (SSTF-BA2201-08).

## Author contributions

H.-W.R. conceived the study. C.-M.Y. led DESTNI development and FACS-based screening, performed proteomics and metabolomics sample preparation and data analysis, and wrote the original draft of the manuscript. J.-Y.J. developed the machine-learning–based identification pipeline for DTB-modified metabolome with the highly sensitive capillary-flow LC-MS system, and generated all DTB-metabolome and proteome LC-MS data with help from J. Kim. C.-R.C. performed LC-MS analyses for initial proof of concept and synthesized DTB–NHS. J. Kang performed AlphaFold2 modeling and molecular docking analyses. S.J. performed cloning and generated stable cell lines. Y.S.P. helped establish the yeast display platform and contributed to DESTNI development. Y.J.C. and Y.P.K. provided metabolite standards and contributed to the construction of the reference spectral library. H.-W.R., J.-S.K. and T.H.Y. supervised the study. All authors reviewed and approved the manuscript.

